# A rarefaction-without-resampling extension of PERMANOVA for testing presence-absence associations in the microbiome

**DOI:** 10.1101/2021.04.06.438671

**Authors:** Yi-Juan Hu, Glen A. Satten

## Abstract

**Background:** PERMANOVA [1] is currently the most commonly used method for testing community-level hypotheses about microbiome associations with covariates of interest. PERMANOVA can test for associations that result from changes in which taxa are present or absent by using the Jaccard or unweighted UniFrac distance. However, such presence-absence analyses face a unique challenge: confounding by library size (total sample read count), which occurs when library size is associated with covariates in the analysis. It is known that *rarefaction* (subsampling to a common library size) controls this bias, but at the potential costs of information loss and the introduction of a stochastic component into the analysis.

**Methods:** Here we develop a non-stochastic approach to PERMANOVA presence-absence analyses that aggregates information over *all* potential rarefaction replicates without actual resampling, when the Jaccard or unweighted UniFrac distance is used. We compare this new approach to three possible ways of aggregating PERMANOVA over multiple rarefactions obtained from resampling: averaging the distance matrix, averaging the (element-wise) squared distance matrix, and averaging the *F*-statistic.

**Results:** Our simulations indicate that our non-stochastic approach is robust to confounding by library size and outperforms each of the stochastic resampling approaches. We also show that, when overdispersion is low, averaging the (element-wise) squared distance outperforms averaging the unsquared distance, currently implemented in the R package `vegan`. We illustrate our methods using an analysis of data on inflammatory bowel disease (IBD) in which samples from case participants have systematically smaller library sizes than samples from control participants.

## Introduction

PERMANOVA [1] is currently the most commonly used method for testing community-level hypotheses about the microbiome. It is used to perform distance-based analyses of taxonomic count data, in order to find associations between community composition and covariates of interest such as environmental factors or clinical outcomes. By choosing an appropriate distance measure, it can be used to analyze either relative abundance data or presence-absence data, in which counts have been converted to a binary variable that indicates whether a taxon is present or absent from a sample. When studying the ecology of the human microbiome and its relationship to disease, PERMANOVA analyses that use relative abundance data appear to be more common. However, many associations between alpha diversity and health have been reported in the medical literature. For example, increased species richness is known to be associated with improved health in the human gut [2], while *decreased* diversity is associated with improved health in the vaginal microbiome [3]. These observations suggest an increased role for presence-absence analyses in human microbiome studies, especially in situations where it is believed that shifts in rare taxa may drive the disease process. In addition, presence-absence analyses may prove to be more robust to the many biases (e.g., extraction and amplification biases) that influence the quantitative results of high-throughput microbiome studies [4].

Presence-absence analyses using PERMANOVA typically adopt either the Jaccard or unweighted UniFrac [5, 6] distance. Presence-absence analyses are particularly sensitive to variability in library size, as the chance of observing rare taxa may depend on sequencing depth. Even though modern 16S rRNA gene sequencing typically yields large library sizes, undersampling of rare taxa may still occur. Without additional information, it is impossible to distinguish between a taxon that is completely absent from the biological specimen, and a taxon that is actually present but has no reads in a particular sample because of under-sampling. For this reason, if library size is associated with any covariate, that covariate may show a spurious association with presence or absence of rare taxa. In theory, confounding by library size may affect the analysis of relative abundance data as well, but the impact is typically much smaller.

Ecologists frequently use rarefaction [7, 8] to overcome the potential confounding effects of library size. Rarefaction corresponds to subsampling reads to a common *rarefaction depth*, which is often the lowest observed library size (after removing outliers). As rarefaction produces data with a single library size, there can be no covariates that are associated with library size. On the other hand, rarefaction can result in a substantial loss of reads and hence statistical power. Multiple rarefaction [8] has been proposed to recover some of the information lost through a single rarefaction. However, rarefaction also introduces a stochastic component to the analysis, and the results obtained even using multiple rarefaction may depend on the number and specifics of the rarefaction replicates sampled.

Here we develop a non-stochastic version of multiple rarefaction that can be used for PERMANOVA presence-absence analyses that use the Jaccard or unweighted UniFrac distance. Our approach is based on an approximation to the result that would be obtained if we could average over *all* potential rarefied datasets. To evaluate the performance of our new approach, we compare it to three possible ways of aggregating the information from multiple rarefied datasets: averaging the distance matrix; averaging the (element-wise) squared distance matrix; and averaging the PERMANOVA *F*-statistic. Only the first strategy, implemented in the `avgdist` function in the `vegan` R package, is widely available, and uses a default value of 100 rarefaction replicates; the averaged distance matrix is then used in a standard PERMANOVA analysis. Although this strategy has already been adopted in several studies [9, 10], its performance has not been compared to the other strategies. In the results section, we present extensive simulation results in which we numerically compared our non-stochastic PERMANOVA to the three resampling strategies; we also evaluate how much of the information lost by rarefaction can be recovered by (an infinite number of) multiple rarefactions. Finally, we apply the methods we discuss to data from a study on inflammatory bowel disease (IBD). We conclude with a discussion section.

## Methods

We briefly review the PERMANOVA approach to establish our notation. Let *X* denote an *n* × *M* design matrix of covariates, where the rows correspond to the *n* samples and the columns correspond to *M* covariates. PERMANOVA can be used to test the effect of a subset of *m* covariates (*m* columns of *X*) using the PERMANOVA *F*-statistic

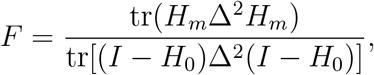

where Δ denotes the distance matrix, Δ^2^ the element-wise squared distance matrix, *H*_*m*_ the hat matrix for the *m* columns we test, *H*_0_ the hat matrix for all columns of *X*, and tr(·) is the trace operation. Note that we drop the multiplier involving the degrees of freedom found in the usual *F*-statistic as PERMANOVA typically assesses significance via permutation.

One way to combine information over *R* rarefaction replicates would be to use the average of the *F*-statistic over the replicates as a test statistic. The significance of the test statistic is assessed by generating the corresponding average of the *F*-statistic calculated from each permutation replicate of data. Even though only Δ varies across rarefaction replicates and can be averaged, averaging the *F* statistic is the “safest” approach since the average over replicates is taken after the final step of the analysis of a single rarefaction replicate. We refer to this approach as PERMANOVA-F(*R*) or F(*R*). A second, easier way to combine information would be to separately average the terms in the numerator and denominator of the *F*-statistic, and then use the ratio of these averages as a test statistic; this corresponds to averaging the *element-wise squared* distance matrices computed for each rarefaction replicate for use in calculating *F*. We refer to this approach as PERMANOVA-D2(*R*) or D2(*R*). A third approach is that implemented in the `avgdist` function in the `vegan` R package, which averages the distance matrices *without squaring* and then uses the average distance to calculate the *F -*statistic. We refer to this approach as PERMANOVA-D(*R*) or D(*R*). PERMANOVA-D2(*R*) and PERMANOVA-D(*R*) only calculate the *F*-statistic once, and are therefore computationally more efficient than PERMANOVA-F(*R*).

PERMANOVA-D(*R*) and PERMANOVA-D2(*R*) calculate numerical approximations to *E*(Δ) and *E*(Δ^2^), respectively, where *E*(·) denotes the expected value over rarefaction replicates. (These expected values are actually conditional on the observed read counts and library sizes; here, we suppress these dependencies for notational convenience.) Non-stochastic versions of these approaches could be provided if these expected values could be calculated analytically. The expectation is taken over the multivariate hypergeometric distribution underlying the generation of a rarefied taxa count table, which is illustrated in Figure 1. Although the ratio form of the Jaccard and unweighted UniFrac distances makes this difficult, it is possible to use the delta method to develop a series approximation to these expected values. We have found that a two-term truncation of this series is easily calculated and provides a very accurate approximation to the exact expected values.

**Figure 1.**
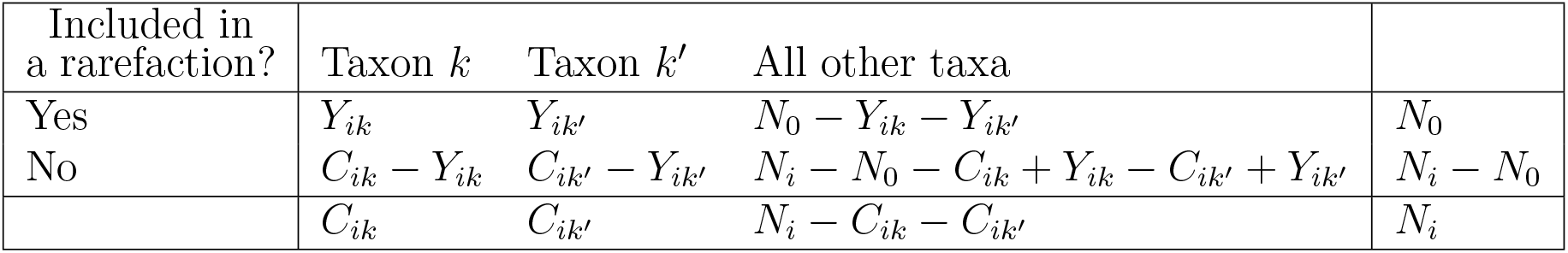
Schematic 2 × 3 table illustrating the multivariate hypergeometric distribution underlying the generation of counts during rarefaction of sample *i. N*_*i*_ is the original library size of sample *i, N*_0_ is the rarefaction depth, *C*_*ik*_ is the original read count of taxon *k* in sample *i*, and *Y*_*ik*_ is the read count of taxon *k* in sample *i* after rarefaction.

More specifically, to develop our method, we first note that both Jaccard and unweighted UniFrac distances between samples *i* and *j* based on a single (rarefied) dataset can be written as

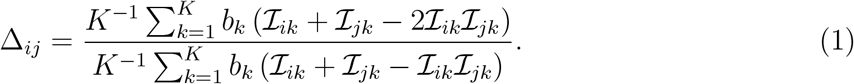

For the Jaccard distance, the summation in (1) is over *K* taxa, the weights *b*_*k*_ = 1 for all *k*, and ℐ_*ik*_ = *I* (*Y*_*ik*_ *>* 0), where *Y*_*ik*_ is the entry in a rarefied taxa count table that corresponds to the *i*th sample and *k*th taxon (see Figure 1). For the unweighted UniFrac, the summation is over the *K* nodes of the phylogenetic tree (excluding the root node) used to calculate the UniFrac distance. For each node *k*, we define 𝒮_*k*_ to be the set of leaf nodes (taxa) that are “below” the *k*th node (i.e., that have node *k* as an ancestor). If we define 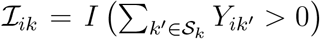 and take weights *b*_*k*_ to be the branch length from node *k* to its parent, then the unweighted UniFrac also has the form given in equation (1).

We define

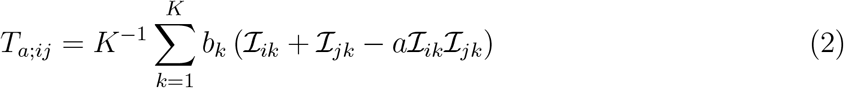

for *a* = 1, 2, and note that Δ_*ij*_ = *T*_2;*ij*_*/T*_1;*ij*_. In the Supplemental Texts, we show using the delta method that we can approximate the expected value of 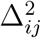 over all rarefaction replicates by

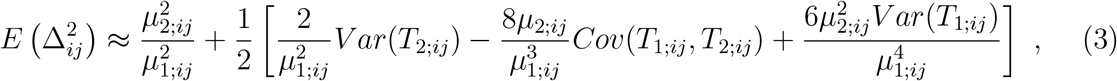

where *µ*_*a*;*ij*_ is the expected value of *T*_*a*;*ij*_ over all rarefaction replicates. Expressions for the moments of *T*_*a*;*ij*_ with respect to the multivariate hypergeometric distribution that are needed to evaluate (3) are also provided in the Supplemental Texts.

We propose to use (3), which is a second-order approximation to 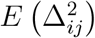, to evaluate the expected values over rarefaction replicates of the numerator and denominator in the PERMANOVA *F*-statistic. Note that the first-order approximation is simply 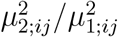, which will also be considered in some of our numerical evaluations. To the extent that (3) provides an accurate evaluation of the expected value of 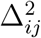, this corresponds to averaging the (element-wise) squared distance matrix over *all* rarefaction replicates; we refer to this approach as PERMANOVA-D2-A_*o*_ or D2-A_*o*_, where *o* denotes the order of the approximation used to calculate 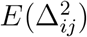.

## Results

### Simulation studies

We created simulated data for 50 cases and 50 controls in which a binary confounder and the case-control status were associated with presence or absence of taxa in the microbiome. We based our simulation on two motivating datasets. One is a microbiome dataset from a smoking study that sequenced the 16S rRNA gene of the throat microbiome [11] and yielded 856 taxa. The other is an ecology dataset from a study of dune meadow vegetation [12] from which the hobby farming subset yielded 21 species (which are also referred to as “taxa” in the sequel for simplicity).

To select taxa that were differentially present in cases and controls for simulations based on the throat microbiome data, we considered two association mechanisms. In the first mechanism (referred to as M1), we randomly selected 100 taxa (after excluding abundant taxa with mean relative abundance *>* 1%), without regard to the phylogeny, to be potentially associated with the case-control status *D*. In the second mechanism (referred to as M2), we selected a cluster of taxa based on the phylogenetic tree to be potentially associated. Specifically, we partitioned the taxa into 20 clusters (lineages) using the partitioning-around-medoids algorithm with pairwise distances given by the between-taxa patristic distance matrix. We selected the largest cluster, which consists of 137 taxa (after excluding abundant taxa with mean relative abundance *>* 1%) to be potentially associated with the case-control status. We independently selected a second set of 100 taxa in M1 and M2 to be associated with a binary confounder *C*, which has a 70% “success” rate in controls but only 30% in cases; note that the sets of taxa associated with *D* and *C* may overlap.

To generate data based on the dune ecology data, we considered only one association mechanism (referred to as M3) in which we randomly selected 5 taxa (after excluding the most abundant taxon) to be potentially associated with the case-control status *D* (which may represent a binary “treatment” variable in an ecology study). The selected taxa had mean relative abundances 0.073, 0.064, 0.039, 0.029, and 0.010. We independently selected a second set of 5 taxa in M3 to be associated with a confounder *C*, which had the same binary structure as in M1 and M2. The 5 selected taxa had mean relative abundances 0.094, 0.040, 0.019, 0.010, and 0.0093. In our simulations, one taxon was common to the sets associated with *D* and *C*.

To simulate data for the *i*th sample, we initially set the relative abundances of all taxa, denoted by *π*_*i*_, to the population mean relative abundances *π* (all entries of which are non-zero), which were estimated from the throat microbiome or dune ecology dataset. For a sample with *D*_*i*_ = 1, we set the entries in *π*_*i*_ to 0, independently for each taxon selected to be associated with *D*, with probability *β* (∈ [0, 1]). We used the value of *β* as a measure of the *effect size*; note that *β* = 0 corresponds to the null hypothesis of no association between the microbiome and case-control status *D*. For a sample with *C*_*i*_ = 1, we further set the entries in *π*_*i*_ to 0, independently for each taxon selected to be associated with *C*, with fixed probability 0.5. To ensure that all entries in *π*_*i*_ sum up to 1, for each sample we increased the probability mass of the most abundant taxon by the total mass that had been set to 0; since the most abundant taxon has sufficient probability mass to ensure it is always present, this operation does not change its presence-absence status. Given *π*_*i*_, we then sampled read count data using the Dirichlet-Multinomial (DM) model. We set the overdispersion parameter to 0.02 in M1 and M2 and 0.0001 in M3, which were estimated from the throat microbiome and dune ecology datasets, respectively. For samples with *D*_*i*_ = 0, we used mean library size 10K in M1 and M2 and 100 in M3. To introduce systematic differences in library size, for samples with *D*_*i*_ = 1, we varied the mean library size from 10K to 5K in M1 and M2 and from 100 to 50 in M3. Given a mean library size *ω*, the library size for each sample was drawn from *N*(*ω, σ* = *ω/*3). Unless otherwise stated, the sampled library size was truncated at 2.5K in M1 and M2 and 20 in M3. The truncation values were used as the rarefaction depth; in this way, no samples were discarded due to rarefaction (corresponding to assuming that samples with problematic library sizes have been removed *a priori* when analyzing real data).

We used the Jaccard distance to analyze data generated using M1 and M3, and the un-weighted UniFrac distance for data generated using M2, corresponding to the most appropriate distance for each scenario. Unless otherwise specified, all calculations were carried out using our implementation of PERMANOVA, `permanovaFL` [13], available in the R package `LDM`. For the new non-stochastic approach, we used the second-order approximation to the expectation of the squared distance unless otherwise stated. In some studies, we included `permanovaFL` applied to the unrarefied distance matrix, referred to as UR. We also evaluated the strategy that simply adjusts the library size as a covariate in `permanovaFL` applied to unrarefied distance matrix, referred to as UR-L. We calculated the type I error (i.e., size) and power for testing the hypothesis of no case-control differences after adjusting for the confounder; the nominal significance level was set to 0.05. Results for size and power were based on 10,000 and 1,000 replicates of data, respectively.

#### Evaluating the approximations to the expected Jaccard and unweighted UniFrac distances

We first evaluated the quality of the first- and second-order approximations to the expectation of the (squared and unsquared) Jaccard and unweighted UniFrac distances, by assessing how many rarefaction replicates were required before we could see evidence that the empirical average over rarefaction replicates was converging to a different limit than our approximation. In Figure 2, we show the log of the maximum percent difference between this empirical average and the first- and second-order approximations, as a function of the log of the number of rarefactions. The linear dependence with slope −1*/*2 is expected given the 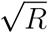 convergence rate for the average over *R* rarefaction replicates, but only when the number of replicates is small enough that the stochastic error in the average is larger than the error in the *o*th (*o* = 1, 2) order approximation. When the number of replicates is large enough that the difference between the average over replicates and the true expected value is small compared to the error in the *o*th order approximation, we expect the empirical log maximum error to switch to a horizontal line as *R* increases. The (approximate) number of replicates at which this change occurs indicates the number of rarefaction replicates that would be required for the average over replicates to obtain a more accurate approximation to 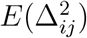 than use of the *o*th order approximation in (3).

**Figure 2.**
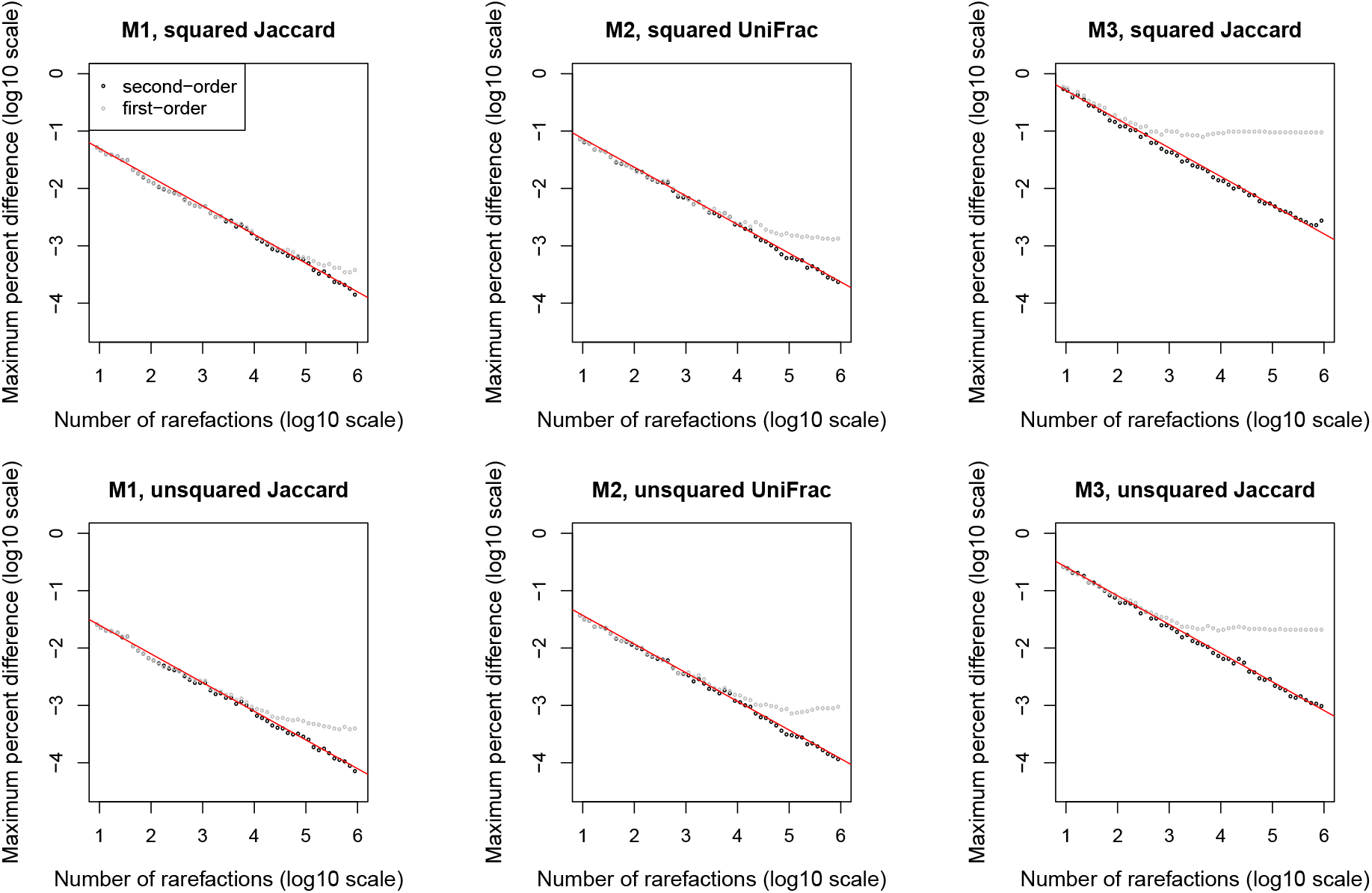
Convergence of the average distance matrix over *R* rarefactions to the first-order (gray) and second-order (black) approximations to the expected distance matrix for three simulated data scenarios. The *y*-coordinate for each point is the maximum over all matrix elements of the absolute value of the percent difference between the average distance matrix and the approximate expected distance matrix, relative to the approximate expected distance matrix. The upper and lower panels correspond to squared and unsquared distance matrices, respectively. The reference line in red is the least-squares fit to the points corresponding to the second-order approximation, with the slope constrained to be −0.5 corresponding to a 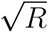 convergence rate. The effect size *β* was set to 0.

For scenarios M1 and M2, we see some evidence that after 10^4^ replicates, the first-order approximation is no longer adequate, while we see little evidence that the second-order approximation is inadequate even when as many as 10^6^ replicates are used. The situation in scenario M3 is different, commensurate with the much smaller number of taxa in this scenario. The first-order approximation is inadequate for as few as 10^2.5^ replicates, while there may be evidence that the empirical average may be outperforming the second-order approximation to the expectation of the squared Jaccard distance when more than 10^6^ replicates are used. Further, the overall error in the empirical average is almost an order of magnitude larger than in scenarios M1 and M2. Taken as a whole, the results in Figure 2 suggest that the second-order approximation performs better than using the empirical average over rarefactions unless the number of rarefactions is extremely large. Even the first-order approximation is better than the empirical average unless the number of taxa is very small.

We performed ordination by first calculating the second-order approximation to the expected (squared) distance matrix, and then projecting off the effect of the confounder *C*, for simulated datasets generated using effect size *β* = 0. Figure 3 shows that this ordination (referred to as “all rarefactions”) is very similar to ordination based on the average (squared) distance matrix over 100 rarefactions in M1 and M2, and moderately similar in M3; this observation is consistent with the higher error rate for scenario M3 seen in Figure 2. As a comparison, we also provided ordination based on unrarefied and singly rarefied distance matrices.

**Figure 3.**
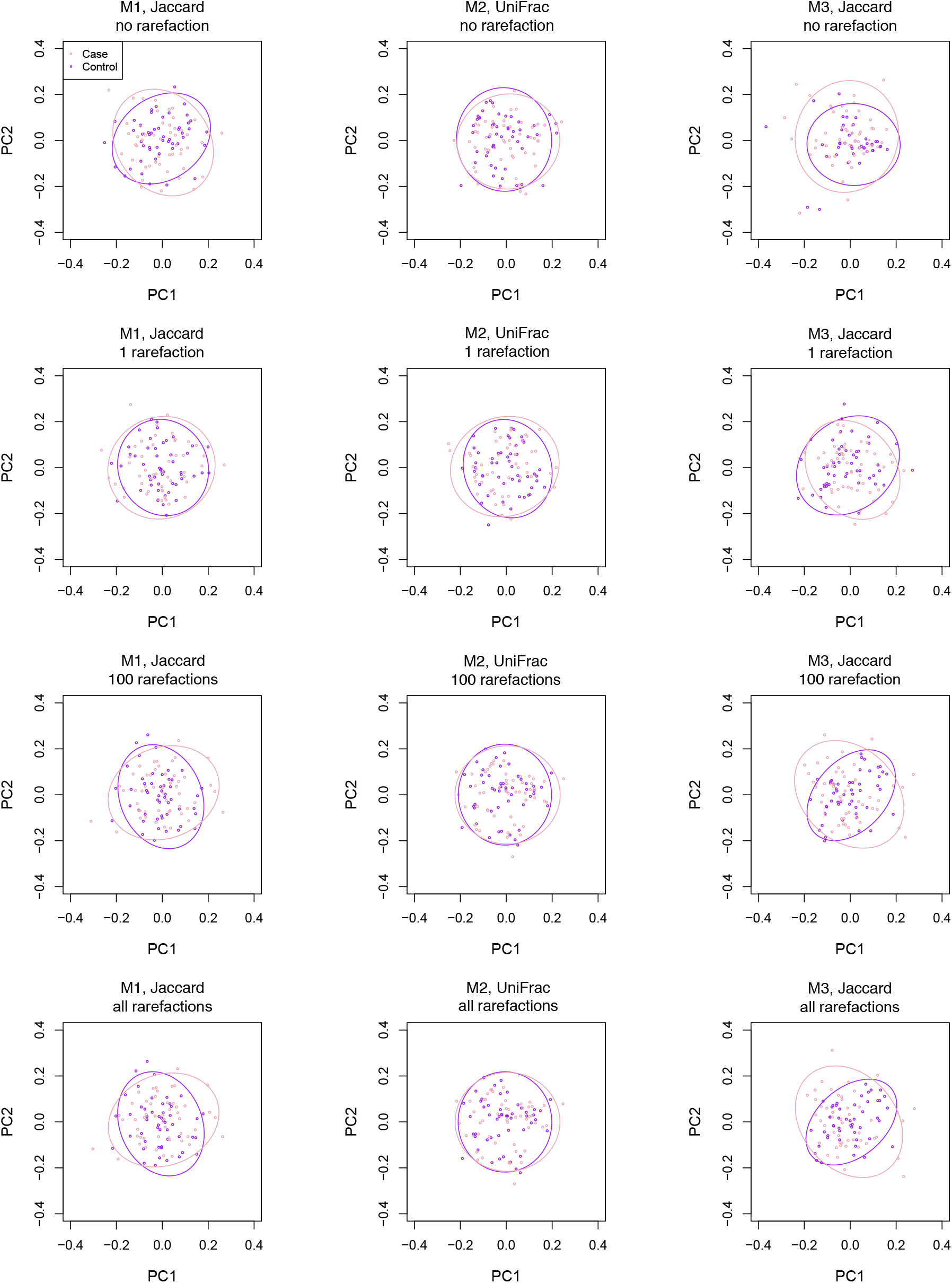
Ordination based on the (squared and centered) distance matrices (after projecting off the effect of the confounder) for the simulated data. The effect size *β* was set to 0. The ellipses are 90% confidence limits.

#### Comparing different versions of PERMANOVA in the presence of systematic differences in library size

We evaluated the size of D2-A_2_, D2(100), F(100), UR, and UR-L. The results are summarized in Figure 4 (upper panel). The proposed methods all control size, regardless of how different the library sizes are distributed between cases and controls. As expected, the size of UR becomes more inflated as the difference in library size increases. It should be noted that UR-L controls size in M1 and M2 and approximately controls size in M3 probably because the library size is essentially a simple binary covariate (i.e., differing in mean in cases and controls); UR-L may not work well in more complex settings if the functional form of the dependence on library size were misspecified. To demonstrate that we have induced substantial confounding effects, we show in Figure S1 that the sizes of D2-A_2_ are as large as 0.2 when the confounder is not controlled for.

**Figure 4.**
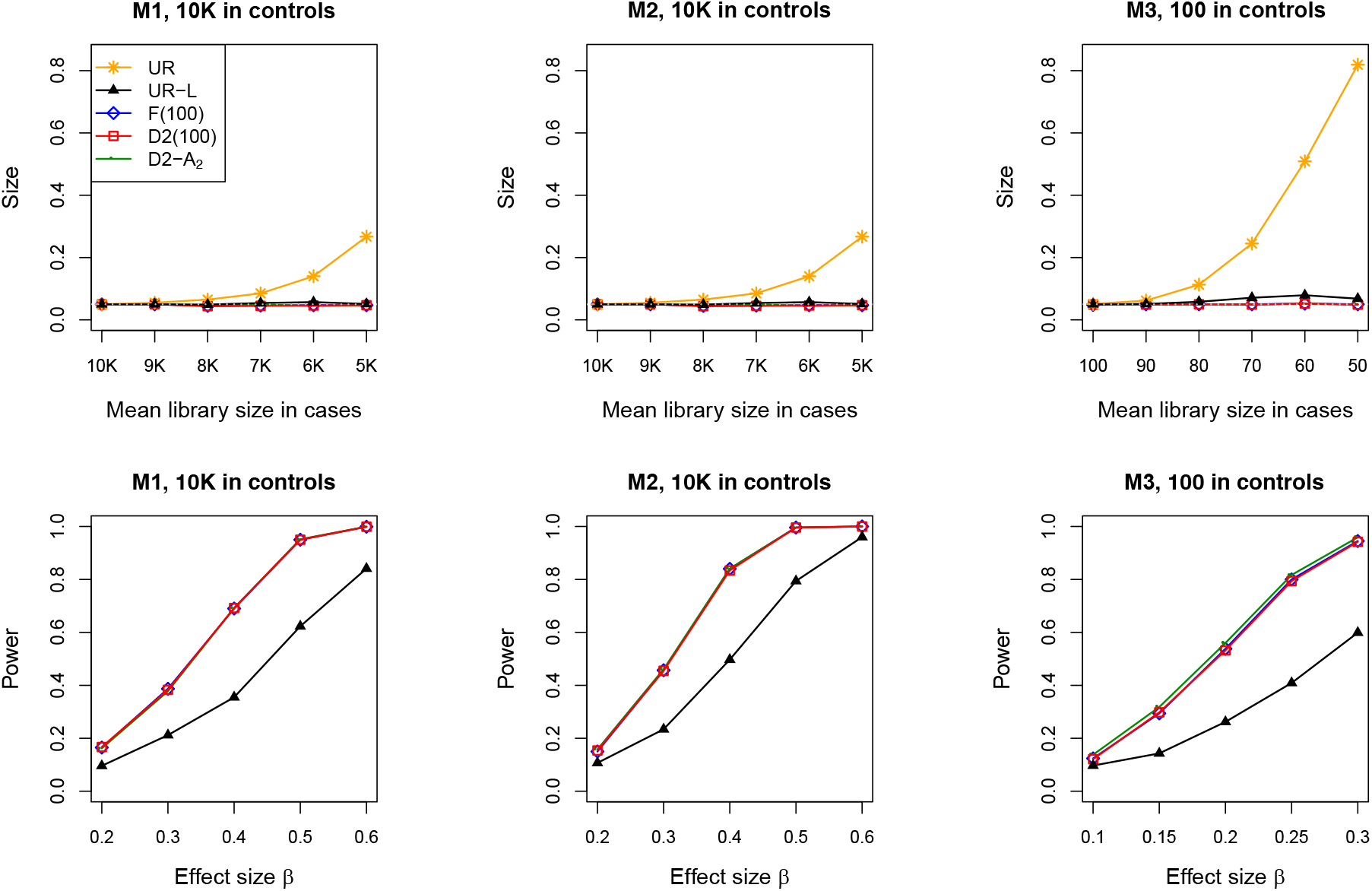
Size and power of the tests, when there are differential library sizes in cases and controls. For assessing power, the mean library sizes are 5K in cases under M1 and M2, and 50 in cases under M3. The gray dotted line represents the nominal significance level of 0.05.

We compared the power of D2-A_2_, D2(100), F(100), and UR-L with a fixed differential library size: 10K and 5K for the mean library sizes of controls and cases, respectively, in M1 and M2, and 100 and 50 in M3. The results are displayed in Figure 4 (lower panel). D2-A_2_ yields the highest power, with D2(100) and F(100) having only slightly less power but UR-L substantially less power. Power results for UR are not shown as this method does not control size. We also compared the power of D2-A_2_ and D2-A_1_ (results not shown). For scenarios M1 and M2, their power was almost identical (less than 0.1% absolute difference); for M3, the power difference was slightly larger, but still less than 0.5%. Figure S2 contrasts the individual *p*-values of D2-A_2_ and D2-A_1_ for 1000 replicates of data. The *p*-values were identical in most cases under M1 and M2 (although differences always favored D2-A_2_); for M3 the differences were larger and more random.

We compared the size and power of D2(*R*) and F(*R*) over different numbers of rarefactions and summarized the results in Figure 5. We added the results of D2-A_2_ as the limit of D2(*R*) when *R* goes to infinity. While all methods control size over all *R*s, aggregating more rarefaction replicates improved power. In M1 and M2 where overdispersion is severe, the bulk of the power increase occurred in the first 5 rarefactions. In M3 where overdispersion is low, power increases were observed even for 100 or more rarefactions. These results indicate the difficulty of choosing *R* in practical settings. In all scenarios, the power of D2(*R*) tracked the power of F(*R*) for all *R*s, suggesting that the ratio of averages is a good approximation to the average of ratios when calculating the PERMANOVA *F*-statistic.

**Figure 5.**
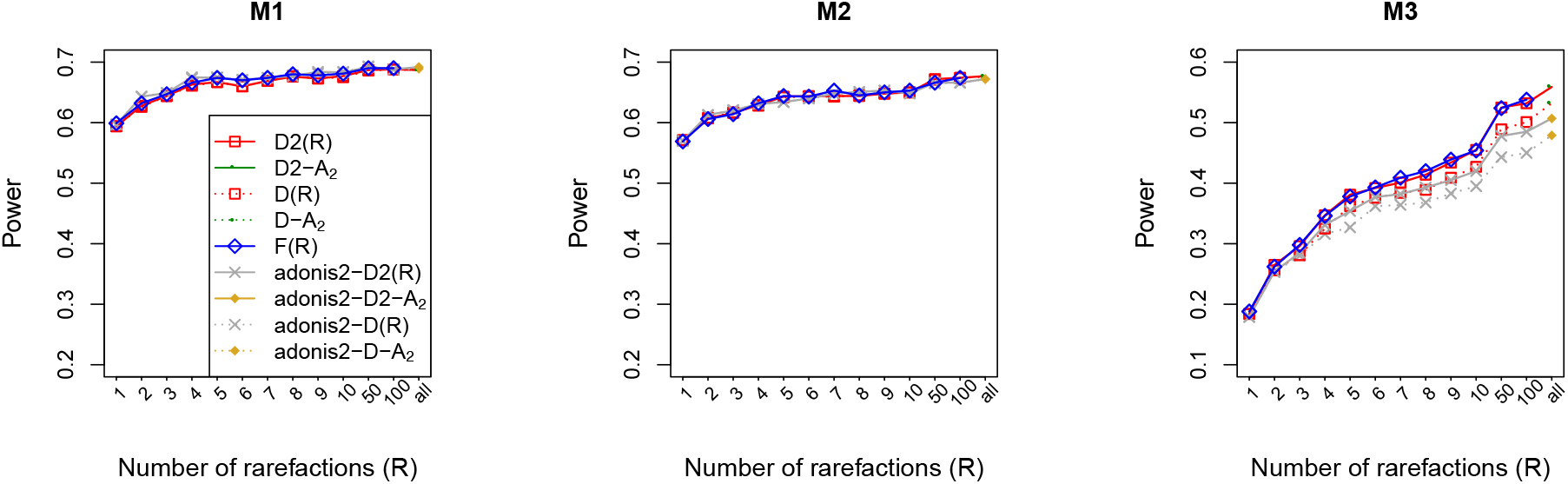
Power of the tests with rarefaction over increasing number of rarefactions. The effect size *β* was set to 0.4, 0.35, and 0.2 under M1, M2, and M3, respectively.

In Figure 5, we also considered whether there was a difference in power between methods that replaced the squared distance matrix used in D2(*R*) and D2-A_2_ by the unsquared distance matrix which we refer to as D(*R*) and D-A_2_. We also included results for methods that replaced the `permanovaFL` function by the `adonis2` function, and refer to them as `adonis2`-D2(*R*), `adonis2`-D2-A_2_, `adonis2`-D(*R*), and `adonis2`-D-A_2_. Note that the existing method implemented in `vegan` with the default value of *R* is `adonis2`-D(100). We demonstrated that, at least in M3, averaging the unsquared distance matrix performs worse than averaging the squared distance matrix, regardless of the number of rarefactions, with either `permanovaFL` or `adonis2`. We also found that the power of `permanovaFL` is higher than the power of `adonis2` in M3. Note that all methods shown in Figure 5 controlled size for all *R*s.

#### The costs of rarefying, when there are no systematic differences in library size

When there is no confounding by library size, the analysis of unrarefied data using UR is valid and can be expected to have optimal power. Then, it is of interest to compare the power of D2-A_2_ to UR. We considered a wide range of the mean library sizes (equal for samples having *D* = 0 and *D* = 1) for scenarios M1, M2, and M3. In each setting, we considered two levels of rarefaction depth, 25% and 10% of the mean library size. Note that on average 90% of the reads are lost in each rarefaction when the rarefaction depth is 10%. The results are displayed in Figure 6. We see that, in M1 and M2 where causal taxa are all rare (i.e., having low relative abundances), rarefaction does lead to loss of power when compared to the analysis of the full data, but the power loss diminishes as the mean library size increases. When the mean library size is large enough, the power of D2-A_2_ eventually reaches that of UR when the rarefaction depth is 25% of the mean. In M3 where several causal taxa are quite abundant, interestingly, D2-A_2_ had higher power than UR. This is not surprising because rarefaction reduces noise at rare taxa that are not associated while having minimal effects on the presence or absence of abundant taxa.

**Figure 6.**
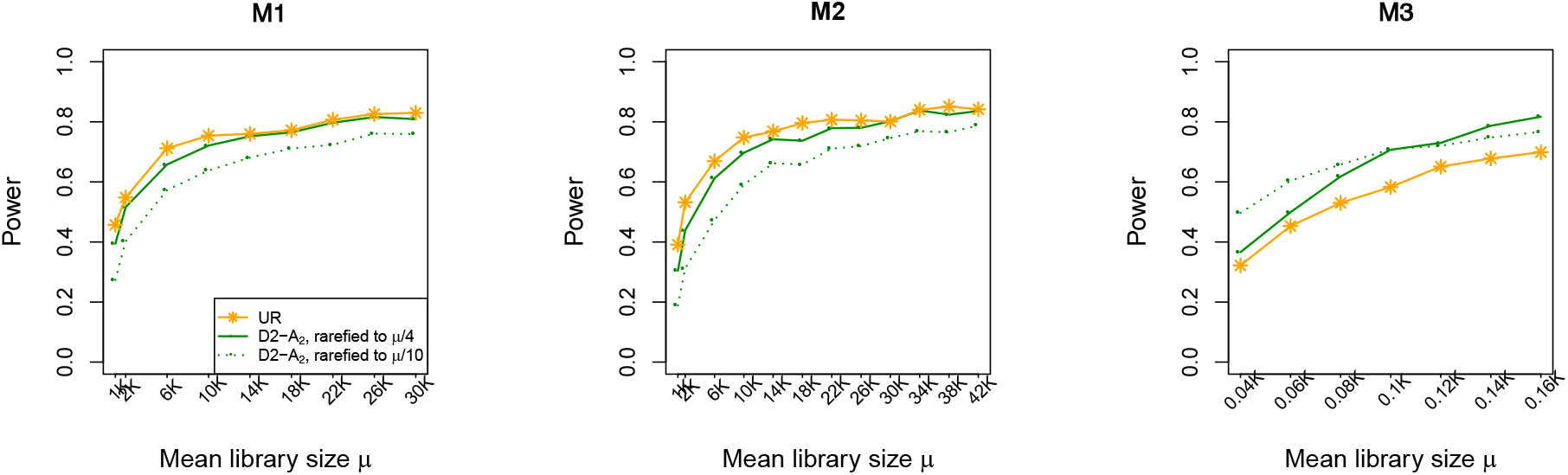
Comparing unrarefied analysis with rarefied analysis, when there are no systematic differences in library size between cases and controls. The effect size *β* was set to 0.4, 0.35, and 0.2 under M1, M2, and M3, respectively.

#### Computation time

The time required to analyze one dataset simulated under M1 using the Jaccard distance was 762 seconds to calculate the average of 100 *F*-statistics for F(100) in both the observed dataset and the 5,000 permutation datasets. It took 8 seconds to calculate the average of 100 squared Jaccard distance matrices used in D2(100) and then an additional 9 seconds to run `permanovaFL` once (with 5,000 permutations). D2-A_*o*_ replaces the average of squared distance in D2(100) by the approximated expectation of squared distance, which took 0.15 or 5 seconds to calculate using the first-order or second-order approximation, respectively. Timings reported here are for a MacBook Pro with 2.9 GHz Intel Core i7 and 16GB memory.

For scenario M2, we used the `GUniFrac` function in the `GUniFrac` R package to calculate the unweighted UniFrac distance for each rarefaction replicate. Analysis of a single dataset using F(100) required 933 seconds, while D2(100) required 99 seconds (of which 90 seconds were used to calculate the average of 100 squared UniFrac distance matrices and 9 seconds for running `permanovaFL`). D2-A_2_ required 673 seconds to calculate the second-order approximation to the expected squared distance, but D2-A_1_ required only 0.5 seconds. Note that by permuting covariates rather than taxa count tables, *E*(Δ) or *E*(Δ^2^) only need to be calculated once, even when significance is determined by permutation.

### Analysis of the IBD data

We analyzed data from case participants diagnosed with Inflammatory Bowel Disease (IBD) and control participants that was first reported by [14]. The goal of our analysis was to test for presence-absence association of the rectal microbiome with IBD status, while controlling for two potential confounders: sex and antibiotic use. Following a common practice, we filtered out taxa that were present in fewer than five samples. We further removed samples with library sizes less than 5,000, which resulted in a loss of 3.4% samples and allowed for a rarefaction depth of 5,221 reads. Our final dataset comprised 284 samples and 2,565 taxa.

Motivated by other researchers [15] who reported substantial confounding of PERMANOVA results due to library size in the same study, we confirmed that the library size distributions were indeed systematically different (Figure 7). Thus, we rarefied the read count data of all 284 samples to the minimum depth 5,221. Similarly to Figure 3, we constructed ordination plots using the squared Jaccard and unweighted UniFrac distances (after removing the effects of sex and antibiotic use [13]) in Figure S3, without rarefaction and with 1, 100, and all rarefactions. These plots demonstrate a clear shift in cases compared with controls both before and after rarefaction. In particular, the ordination based on our calculated expected distance matrix (i.e., all rarefactions) is almost identical to the ordination based on the average distance matrix of 100 rarefactions and even similar to the ordination based on one rarefaction; all the rarefaction-based ordinations show some differences from ordinations based on the unrarefied data.

**Figure 7.**
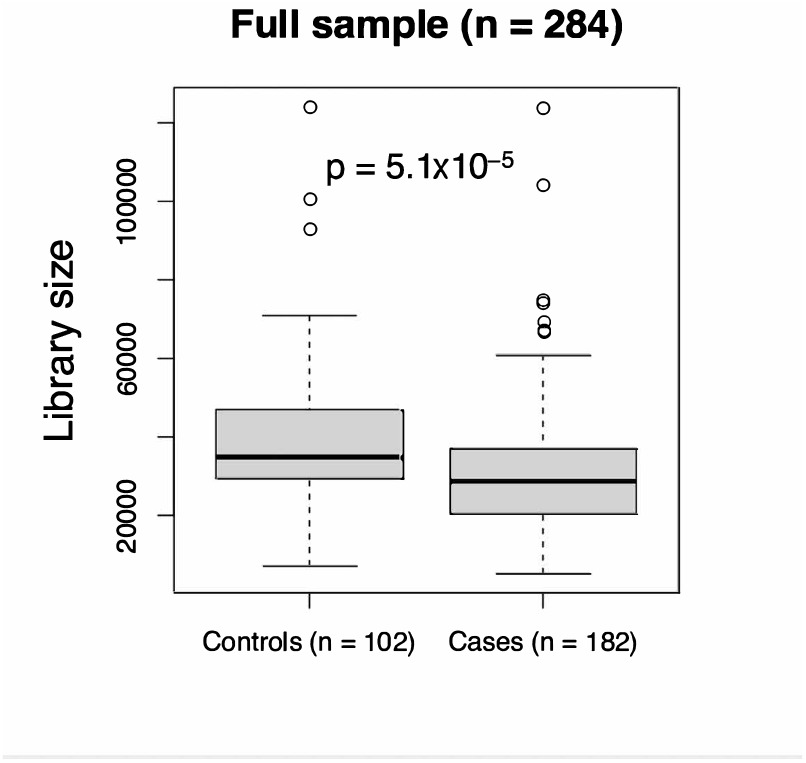
Differential distributions of library sizes in cases and controls of the IBD data. The *p*-value was generated by the Wilcoxon rank-sum test.

We applied F(*R*) and D2(*R*) with *R* increasing from 1 to 100 and D2-A_2_, using both the Jaccard and unweighted UniFrac distances. The *p*-values of all tests as shown in Figure 8 (upper) are equal to the minimum possible value (i.e., 1*/*5001 where we chose 5,000 to be the maximum number of permutations), regardless of the number of rarefactions. These *p*-values indicated a very strong presence-absence association of the IBD status with the rectal microbiome at the community level, which is not surprising given the clear shift of case ordinations relative to controls even with only one rarefaction. In Figure 8, we plot results from D2-A_2_ as the limiting value of D2(*R*) obtained as *R* increases.

**Figure 8.**
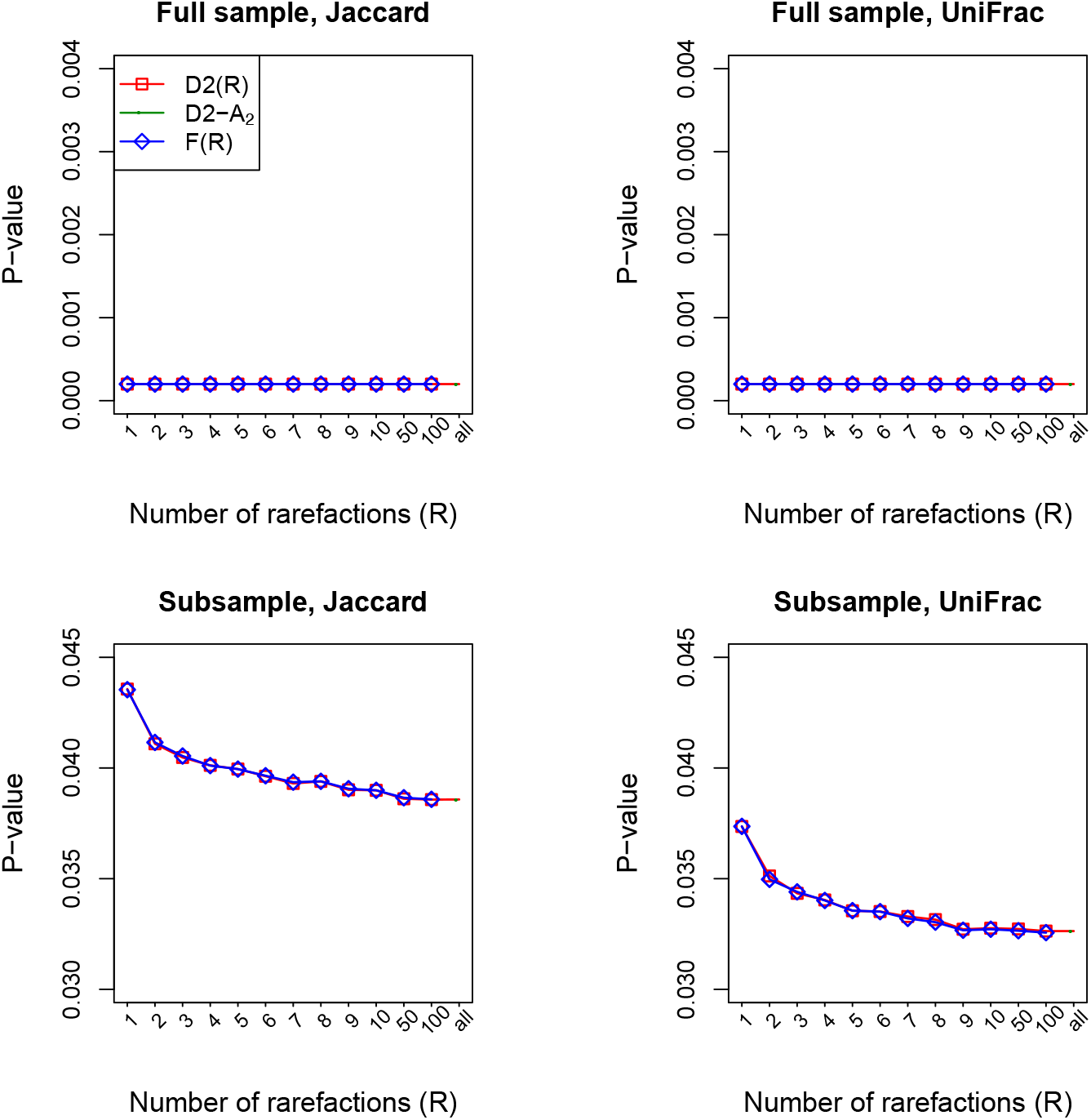
*P* -values for testing presence-absence association of the rectal microbiome with the IBD status. The *p*-values in the lower panel are average *p*-values from 1000 subsets of the original data, each consisting of 50 randomly selected samples.

Because the association was so strong in the IBD data, we analyzed subsets of data, each consisting of 50 randomly selected samples. We obtained *p*-values using F(*R*), D2(*R*), and D2-A_2_ for each of 1000 such subsets, and report the average *p*-values. The average *p*-values obtained using F(*R*) and D2(*R*) are in close agreement for all values of *R*. As *R* increases, *p*-values using F(*R*) and D2(*R*) decrease and eventually converge to the *p*-value obtained using D2-A_2_. These patterns are consistent with our observations using simulated data, reported in Figure 5. While the results of F(*R*) and D2(*R*) depend on the value of *R*, D2-A_2_ produces a single value which is, additionally, the lowest *p*-value among the three methods on average. Thus, we conclude that D2-A_2_ provides the best presence-absence analyses of these data.

## Discussion

We have presented a non-stochastic method for a rarefaction-based analysis of presence-absence data using PERMANOVA. This approach uses the delta method to approximate the analytic average of the squared distance matrix over all rarefaction replicates. We showed that our non-stochastic method has higher power than each of the three stochastic approaches to aggregating information over multiple rarefaction replicates. Further, our analytical average of the squared distance matrix can be used as if it were an ordinary distance matrix, for example, for performing ordination or constructing an omnibus test that combines results from multiple distances [16]. We have included all the methods described here in our `LDM` R package for microbiome data analysis, which is available on GitHub at https://github.com/yijuanhu/LDM.

We have previously developed the linear decomposition model (LDM) that unifies the community-level and taxon-level tests into one framework [13]. We also extended the LDM for testing presence-absence associations based on multiple rarefactions [17]; some of those results mirror the findings we obtained here. Specifically, we compared a stochastic approach that averages the LDM *F*-statistic from *R* rarefaction replicates to a non-stochastic approach that separately calculates the expected values of the residual sum-of-squares (RSS) terms in the numerator and denominator of the LDM *F*-statistic with respect to the rarefaction operation. Parallel to the results we found here, the non-stochastic approach had superior performance. Unlike the case of PERMANOVA, the expected values over rarefaction replicates for the LDM could be calculated *exactly*.

When the distributions of library size are similar between cases and controls, the relative power of the unrarefied (UR) and rarefied analysis (D2-A_2_) depends on whether rare or abundant taxa are involved in the association. Although unrarefied analysis typically gives the best power when only rare taxa are associated, the power lost by rarefaction is minimal, as long as the overall sequencing depth is moderately high and the rarefaction depth is not too low. When abundant taxa are associated, rarefaction can even improve power. Given the robustness of rarefied analysis to any systematic variation in library size, it seems prudent to use D2-A_2_ even when there is no obvious difference in the distributions of library size.

The efficiency of rarefied analysis depends on the rarefaction depth. In general, the guideline for choosing a rarefaction depth is similar to other applications that use rarefaction: the depth should be low enough to include most samples, but can exclude samples with anomalously low library sizes. We also saw that the power of D2-A_2_ could be slightly higher than that of D2-A_1_, with the largest differences occurring when the number of taxa is small. Thus we recommend use of D2-A_2_ whenever it is computationally feasible; fortunately the situations in which D2-A_2_ is time-consuming to calculate are exactly the situations for which the performance of D2-A_1_ is nearly identical to D2-A_2_. When the number of taxa is small, the second-order correction D2-A_2_ should always be used. Finally, since PERMANOVA typically permutes covariates rather than residuals, we also note that the second-order approximation only needs to be calculated once for each analysis.

## Conclusions

We present a non-stochastic approach to PERMANOVA presence-absence analyses that aggregates information over *all* potential rarefaction replicates without actual resampling, when the Jaccard or unweighted UniFrac distance is used. Our approach is robust to confounding by library size and outperforms the three possible stochastic resampling approaches we have considered. As PERMANOVA is currently the most commonly used method for testing community-level hypotheses about the microbiome, our extension of PERMANOVA for robust and powerful analysis of presence-absence associations represents substantial advances of PERMANOVA to be better used to advance scientific knowledge.

## Availability of data and materials

The R package `LDM` is available on GitHub at https://github.com/yijuanhu/LDM in formats appropriate for Macintosh, Linux, or Windows systems.

## Authors’ contributions

YJH conceived the study, developed the method, performed simulation studies, analyzed the data, and wrote the manuscript. GAS conceived the study, developed the method, and wrote the manuscript. Both authors read and approved the final manuscript.

## Consent for publications

No consent for publication was required for this study; all microbiome datasets used here are publicly available.

## Ethics approval and consent to participate

This study only involved secondary analyses of existing, de-identified datasets; as such it does not require separate IRB consent.

## Acknowledgements

Not applicable.

