## Supplemental Figures and Texts for "A rarefaction-without-resampling extension of PERMANOVA for testing presence-absence associations in the microbiome"

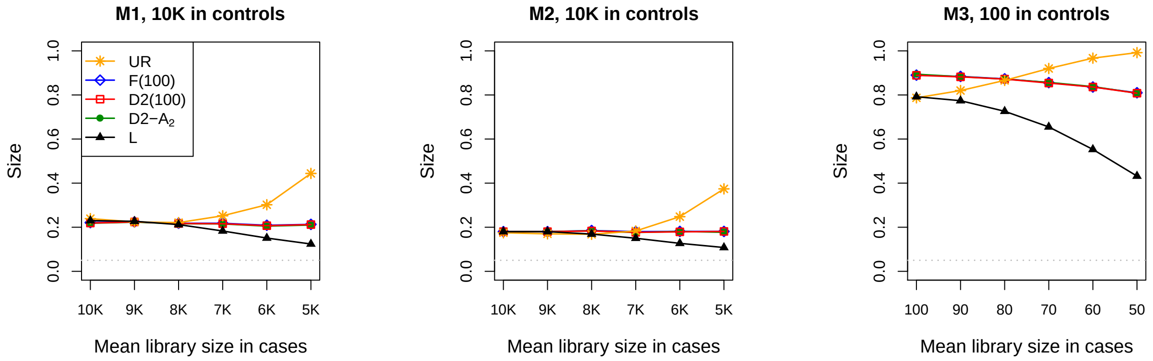


**Figure S1.** Size of the tests, when the confounder is not controlled for. The gray dotted line represents the nominal significance level of 0.05.

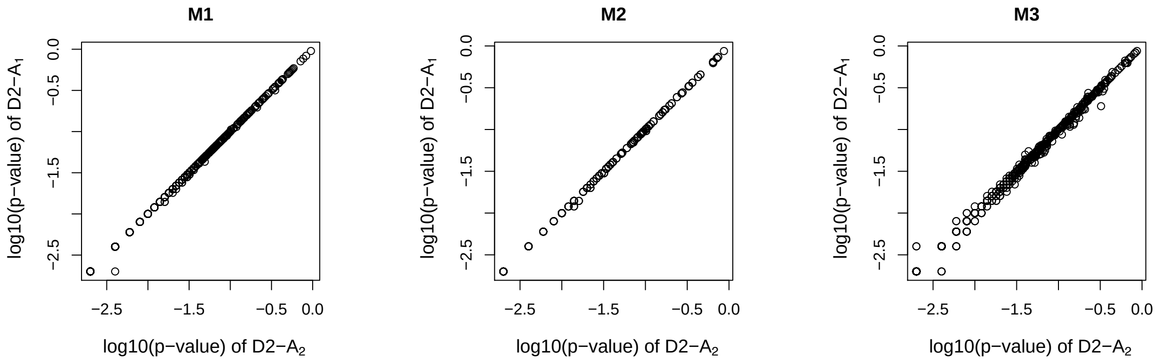


**Figure S2.** Contrasting $p$-values generated by D2-$A_2$ and D2-$A_1$ for 1000 replicates of data. The mean library sizes are 10K and 5K in controls and cases, respectively, under M1 and M2, and 100 and 50 in controls and controls under M3. The effect size $\beta$ was set to 0.4 under M1 and M2 and 0.2 under M3.

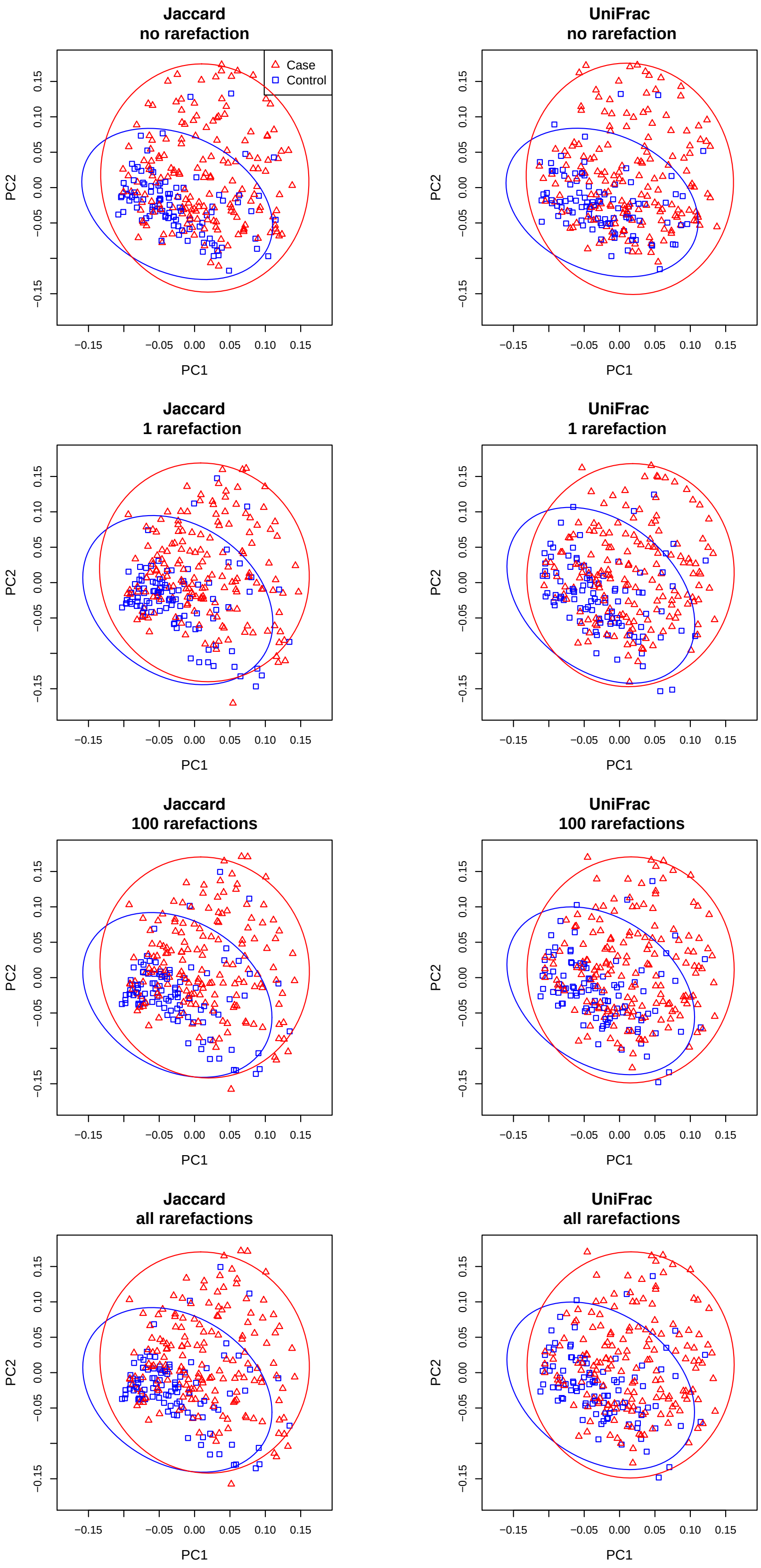


**Figure S3.** Ordination based on the (squared and centered) Jaccard and unweighted UniFrac distance matrices (after projecting off sex and antibiotic use variables) for the IBD data. The ellipses are 95% confidence limits for case and control clusters.

### Supplemental Texts: Calculating the Expected Jaccard and Unweighted UniFrac Distances

#### S1. Statistics of Rarefaction

To calculate the expectation of the squared Jaccard distance, we will need the following quantities. From the multivariate hypergeometric (MH) distribution illustrated in Figure 1, we have

$$\Pr\Big(Y_{ik}=0\Big) = \frac{\binom{C_{ik}}{0}\binom{N_i - C_{ik}}{N_0}}{\binom{N_i}{N_0}} := \overline{p}_{ik} := 1 - p_{ik},$$

and

$$\Pr\Big(Y_{ik}=0 \ \& \ Y_{ik'}=0\Big) = \frac{\binom{C_{ik}}{0}\binom{C_{ik'}}{0}\binom{N_i - C_{ik} - C_{ik'}}{N_0}}{\binom{N_i}{N_0}} := \overline{P}_{i;kk'},$$

where := is used to define a new quantity. In terms of these quantities, it is easy to show that

$$\Pr\Big(Y_{ik}=1 \ \& \ Y_{ik'}=1\Big) := P_{i;kk'} = p_{ik} + p_{ik'} + \overline{P}_{i;kk'} - 1.$$

Since $\mathcal{I}_{ik} = I\left(Y_{ik} > 0\right)$, we have $E\left(\mathcal{I}_{ik}\right) = p_{ik}$ and $E(\mathcal{I}_{ik}\mathcal{I}_{ik'}) = P_{i;kk'}$.

For the unweighted UniFrac distance, we define $\mathcal{I}_{ik} = I\left(\sum_{k'\in\mathcal{S}_k} Y_{ik'} > 0\right)$. We have from the collapsibility of the MH distribution that

$$\Pr\Big(\mathcal{I}_{ik}=0\Big) = \Pr\Big(\sum_{k'\in\mathcal{S}_k} Y_{ik'} = 0\Big) = \frac{\binom{N_i - \sum_{k'\in\mathcal{S}_k} C_{ik'}}{N_0}}{\binom{N_i}{N_0}} := \overline{p}_{ik} := 1 - p_{ik}.$$

Similar to the Jaccard case, let

$$\Pr\Big(\mathcal{I}_{ik}=0 \ \& \ \mathcal{I}_{ik'}=0\Big) = \Pr\left(\sum_{k'\in\mathcal{S}_k\cup S_{k'}} Y_{ik'} = 0\right) := \overline{P}_{i;kk'}.$$

Then, as with the Jaccard distance, we have

$$\Pr\Big(\mathcal{I}_{ik}=1 \ \& \ \mathcal{I}_{ik'}=1\Big) := P_{i;kk'} = p_{ik} + p_{ik'} + \overline{P}_{i;kk'} - 1,$$

and so $E\left(\mathcal{I}_{ik}\right) = p_{ik}$ and $E(\mathcal{I}_{ik}\mathcal{I}_{ik'}) = P_{i;kk'}$. Note that the precise definitions of $\mathcal{I}_{ik}$, $p_{ik}$ and $P_{i;kk'}$ are different for the Jaccard and unweighted UniFrac distances.

### S2. Delta-Method Approximation to the Expected Jaccard and Unweighted UniFrac Distances

We use the Delta method to derive a second-order approximation to $E(\Delta^2)$, as well as to $E(\Delta)$ which is also included in our numerical evaluation. For simplicity we drop the indices $i, j$. If we expand $\Delta = T_2/T_1$ for $T_1$ around $\mu_1$ and $T_2$ around $\mu_2$ to second order, we obtain

$$E(\Delta) \approx \frac{\mu_2}{\mu_1} + \frac{1}{2}\left[-\frac{2}{\mu_1^2}Cov(T_1, T_2) + \frac{2\mu_2 Var(T_1)}{\mu_1^3}\right] = \frac{\mu_2}{\mu_1} + \frac{\mu_2 m_{11} - \mu_1 m_{12}}{\mu_1^3}$$

where

$$m_{ab} = E(T_a T_b).$$

Similarly, if we expand $\Delta^2 = T_2^2/T_1^2$ for $T_1$ around $\mu_1$ and $T_2$ around $\mu_2$ to second order, we obtain

$$E(\Delta^2) \approx \frac{\mu_2^2}{\mu_1^2} + \frac{\mu_1^2 m_{22} - 4\mu_1\mu_2 m_{12} + 3\mu_2^2 m_{11}}{\mu_1^4}.$$

To justify use of the delta method, recall from equation (2) that $T_1$ and $T_2$ can be considered as means over the stochastic process that generates data at a random taxon (for the Jaccard distance) or a random branch (for the unweighted UniFrac distance). Thus, we can expect that the second moments are of order $\mathcal{O}(K^{-1})$ and that each successive moment will be smaller by a factor of $K^{-\frac{1}{2}}$.

### S3. Calculation of Moments

We show the details for the UniFrac distance; the results for the Jaccard distance are easily obtained from those of the UniFrac by setting all weights $b_k = 1$ and choosing the sum over $k$ to index taxa rather than nodes in the phylogenetic tree. The first moments are easy, and we find

$$\mu_a = E\left\{\sum_{k=1}^{K} b_k\Big(\mathcal{I}_{ik} + \mathcal{I}_{jk} - a\mathcal{I}_{ik}\mathcal{I}_{jk}\Big)\right\} = \sum_k b_k\Big(p_{ik}+p_{jk}-ap_{ik}p_{jk}\Big) = p_{i\cdot}^{(1)}+p_{j\cdot}^{(1)}-a\sum_k b_k p_{ik}p_{jk},$$

where we define $p_{i\cdot}^{(1)} = \sum_{k=1}^K b_k p_{ik}$. The second moment requires some work and we first expand

$$\begin{aligned} m_{a_1a_2} &= E\left\{\sum_{k=1}^K\sum_{k'=1}^K b_kb_{k'}\Big(\mathcal{I}_{ik}+\mathcal{I}_{jk}-a_1\mathcal{I}_{ik}\mathcal{I}_{jk}\Big)\Big(\mathcal{I}_{ik'}+\mathcal{I}_{jk'}-a_2\mathcal{I}_{ik'}\mathcal{I}_{jk'}\Big)\right\} \\ &= E\left\{\sum_{k=1}^K b_k^2\Big(\mathcal{I}_{ik}+\mathcal{I}_{jk}-a_1\mathcal{I}_{ik}\mathcal{I}_{jk}\Big)\Big(\mathcal{I}_{ik}+\mathcal{I}_{jk}-a_2\mathcal{I}_{ik}\mathcal{I}_{jk}\Big)\right\} \qquad \text{(S1)} \\ &\quad + E\left\{\sum_{k,k'=1,k\neq k'}^K b_kb_{k'}\Big(\mathcal{I}_{ik}+\mathcal{I}_{jk}-a_1\mathcal{I}_{ik}\mathcal{I}_{jk}\Big)\Big(\mathcal{I}_{ik'}+\mathcal{I}_{jk'}-a_2\mathcal{I}_{ik'}\mathcal{I}_{jk'}\Big)\right\}. \qquad \text{(S2)} \end{aligned}$$

The first term (S1) can be factored out to give, by noting that $\mathcal{I}_{ik}^2 = \mathcal{I}_{ik}$,

$$\begin{aligned} & E\left\{\sum_{k=1}^K b_k^2\Big[\mathcal{I}_{ik}+\mathcal{I}_{jk}+2\mathcal{I}_{ik}\mathcal{I}_{jk}-2(a_1+a_2)\mathcal{I}_{ik}\mathcal{I}_{jk}+a_1a_2\mathcal{I}_{ik}\mathcal{I}_{jk}\Big]\right\} \\ &= \sum_{k=1}^K b_k^2\left\{p_{ik}+p_{jk}+\Big[2-2(a_1+a_2)+a_1a_2\Big]p_{ik}p_{jk}\right\} \\ &= p_{i\cdot}^{(2)}+p_{j\cdot}^{(2)}+\Big[2-2(a_1+a_2)+a_1a_2\Big]\sum_{k=1}^K b_k^2p_{ik}p_{jk}, \end{aligned}$$

where we define $p_{i\cdot}^{(2)} = \sum_{k=1}^K b_k^2 p_{ik}$. The second term (S2) can be factored out to give

$$\begin{aligned} &\sum_{k,k'=1,k\neq k'}^K b_kb_{k'}\Big\{E\Big(\mathcal{I}_{ik}\mathcal{I}_{ik'}+\mathcal{I}_{ik}\mathcal{I}_{jk'}+\mathcal{I}_{ik'}\mathcal{I}_{jk}+\mathcal{I}_{jk}\mathcal{I}_{jk'}\Big) \\ &-E\Big(a_2\mathcal{I}_{ik}\mathcal{I}_{ik'}\mathcal{I}_{jk'}+a_2\mathcal{I}_{jk}\mathcal{I}_{ik'}\mathcal{I}_{jk'}+a_1\mathcal{I}_{ik'}\mathcal{I}_{ik}\mathcal{I}_{jk}+a_1\mathcal{I}_{jk'}\mathcal{I}_{ik}\mathcal{I}_{jk}\Big)+a_1a_2E\Big(\mathcal{I}_{ik}\mathcal{I}_{ik'}\mathcal{I}_{jk}\mathcal{I}_{jk'}\Big)\Big\} \\ &= \sum_{k,k'=1,k\neq k'}^K b_kb_{k'}\Big\{P_{i;kk'}+2p_{ik}p_{jk'}+P_{j;kk'}-\Big(a_2p_{jk'}+a_1p_{jk}\Big)P_{i;kk'}-\Big(a_2p_{ik'}+a_1p_{ik}\Big)P_{j;kk'}+a_1a_2P_{i;kk'}P_{j;kk'}\Big\}. \end{aligned}$$

The term that involves $p_{ik}p_{jk'}$ can be simplified by noting that

$$2\sum_{k,k'=1,k\neq k'}^K b_kb_{k'}\,p_{ik}p_{jk'} = 2\sum_{k=1}^K\sum_{k'=1}^K b_kb_{k'}p_{ik}p_{jk'}-2\sum_{k=1}^K b_k^2p_{ik}p_{jk} = 2p_{i\cdot}^{(1)}p_{j\cdot}^{(1)}-2\sum_{k=1}^K b_k^2p_{ik}p_{jk}.$$

Combining the results of the two terms (S1) and (S2), we have the second moment to be

$$\begin{aligned} m_{a_1a_2} &= p_{i\cdot}^{(2)}+p_{j\cdot}^{(2)}+2p_{i\cdot}^{(1)}p_{j\cdot}^{(1)}+\Big[a_1a_2-2(a_1+a_2)\Big]\sum_{k=1}^K b_k^2p_{ik}p_{jk} \\ &\quad+\sum_{k,k'=1,k\neq k'}^K b_kb_{k'}\Big\{\Big(1-a_1p_{jk}-a_2p_{jk'}\Big)P_{i;kk'}+\Big(1-a_1p_{ik}-a_2p_{ik'}\Big)P_{j;kk'}+a_1a_2P_{i;kk'}P_{j;kk'}\Big\}. \end{aligned}$$

To speed up computation of the three terms in $m_{a_1a_2}$ that involve $\sum_{k,k'=1,k\neq k'}^{K}$, we modify the sum over $(k,k')$ to only consider $k<k'$ values; this will allow us to cut the number of terms in the sum by a factor of 2. This requires a bit of care, since the case $a_1 \neq a_2$ has some asymmetry, so we cannot just impose the restriction and multiply by a factor of 2 to compensate. To start this, note that $\sum_{k,k'=1,k\neq k'}^{K} = \sum_{k=1}^{K-1}\sum_{k'=k+1}^{K} + \sum_{k=2}^{K}\sum_{k'=1}^{k-1}$. For the first term involving $\sum_{k,k'=1,k\neq k'}^{K}$, we define the first double summation to be

$$\phi_{ij}(a_1,a_2) = \sum_{k=1}^{K-1}\sum_{k'=k+1}^{K} b_k b_{k'}\Big(1 - a_1 p_{jk} - a_2 p_{jk'}\Big) P_{i;kk'}.$$

Then the second double summation can be written as

$$\sum_{k=2}^{K}\sum_{k'=1}^{k-1} b_k b_{k'}\Big(1 - a_1 p_{jk} - a_2 p_{jk'}\Big) P_{i;kk'} = \sum_{k'=1}^{K-1}\sum_{k=k'+1}^{K} b_k b_{k'}\Big(1 - a_1 p_{jk} - a_2 p_{jk'}\Big) P_{i;kk'} = \phi_{ij}(a_2,a_1),$$

where the first equation follows from exchanging the order of the sums. Thus, the first term involving $\sum_{k,k'=1,k\neq k'}^{K}$ is $\phi_{ij}(a_1,a_2) + \phi_{ij}(a_2,a_1)$. The second term involving $\sum_{k,k'=1,k\neq k'}^{K}$ only differs by exchanging $i$ and $j$, and gives $\phi_{ji}(a_1,a_2) + \phi_{ji}(a_2,a_1)$. The situation with the third term is easier, as the summation does not involve $a_1$ and $a_2$:

$$a_1 a_2 \sum_{k,k'=1,k\neq k'}^{K} b_k b_{k'} P_{i;kk'} P_{j;kk'} = 2 a_1 a_2 \sum_{k=1}^{K-1}\sum_{k'=k+1}^{K} b_k b_{k'} P_{i;kk'} P_{j;kk'}.$$

The calculation for the Jaccard distance, obtained by setting $b_k = 1$ for any $k$, is nearly identical. The only difference is that with the weights removed, we have $p_{i\cdot}^{(1)} = p_{i\cdot}^{(2)}$.
